# A Rapid, Cost-Effective Method for High-Yield DNA Purification from PCR Products and Agarose Gels

**DOI:** 10.1101/2024.11.13.623516

**Authors:** Susheel Fatima, Maliha Javed, Hijab Zahra, Irfan Hussain, Afsar Ali Mian

**Affiliations:** Centre for Regenerative Medicine and Stem Cells Research, The Aga Khan University, Karachi, Pakistan

## Abstract

Nucleic acid purification is essential for molecular biology workflows, enabling successful downstream applications like cloning, sequencing, and PCR amplification. While commercial kits are widely used for DNA extraction from PCR products and agarose gels, their high costs can burden resource-limited laboratories. To provide a viable alternative, we developed an optimized, cost-effective in-house protocol for high-yield DNA purification. This study evaluates the performance of our protocol for plasmid DNA and PCR products, comparing it with commercial kits from Qiagen, Thermo Fisher, and WizBio in terms of cost, time, DNA concentration, and purity. Results from gel electrophoresis demonstrated that the in-house protocol significantly enhances PCR product clarity and reduces background smearing, yielding high-purity DNA compatible with sensitive applications. Restriction-digested plasmid samples showed successful ligation and transformation in E. coli (DH5α), with Sanger sequencing chromatogram further confirming the sequence integrity of the purified DNA. Our findings highlight the in-house protocol as a cost-effective, efficient, and reliable alternative to commercial kits, delivering high-quality DNA suitable for various molecular applications. This method offers an accessible and practical solution for laboratories seeking to optimize DNA purification under budget constraints.

## Introduction

Efficient purification of PCR products and DNA fragments from agarose gels is essential in molecular biology(Aldrich and Cullis 1993), enabling successful downstream applications such as sequencing(Maurisse et al. 2006), restriction digestion(Diogo et al. 2000), cloning(Ferreira et al. 2023), labeling(Vomelová et al. 2009), ligation(Fu et al. 2021), in vitro transcription(Carapito et al. 2023), and in situ hybridization(Vomelová et al. 2009). High DNA yield and purity are critical for these applications, but conventional purification methods often rely on silica-based DNA binding in the presence of chaotropic salts(Maurisse et al. 2006; Whitlock et al. 2008). While effective, this approach is associated with high costs, extended processing times, and limited DNA yield, especially in laboratories with budget constraints or high throughput demands(Stadler et al. 2004).

A survey of the literature and insights from research discussions suggest that cost and time inefficiencies are significant challenges for many laboratories using commercial DNA extraction kits. Proprietary reagents, low yields for small DNA fragments, and the need for specialized equipment further hinder accessibility, flexibility, and overall laboratory efficiency(Ahmed et al. 2013). These limitations disproportionately impact resource-limited labs, which may find commercial options prohibitively expensive and restrictive for routine DNA purification(Depristo et al. 2011).

To address these challenges, we developed an optimized in-house DNA purification protocol tailored for agarose gel and PCR product extraction. Our protocol incorporates modifications to traditional methods that enhance DNA recovery, reduce cost, and minimize processing time, providing a practical and accessible alternative to commercial kits. By eliminating costly reagents, this in-house method achieves high yield and purity at a fraction of the cost, making it well-suited for PCR amplification, cloning, and sequencing. Key benefits of this protocol include its simplicity, rapid execution, and cost-effectiveness, positioning it as a valuable solution for molecular workflows requiring high-quality DNA. Furthermore, the protocol demonstrates consistent extraction efficiency across various DNA templates, underscoring its broad applicability and utility for modern molecular biology labs. As the demand for reliable and affordable DNA purification grows, this optimized protocol offers a timely advancement, enabling researchers to enhance their workflows while maintaining rigorous standards of DNA quality.

## Methods

### PCR Amplification, Primer Design, and Restriction Digestion

To facilitate the efficient extraction of PCR products and DNA fragments, PCR amplification was performed using specifically designed primers to target distinct DNA regions, producing amplicons suitable for downstream applications. The details of the primers used were given in **Table 1**. The primers were optimized for specificity and amplification efficiency, with PCR conditions adjusted to produce high-yield, specific amplicons. The sanger sequencing were done commercially from Eurofins.

**Table 1:**
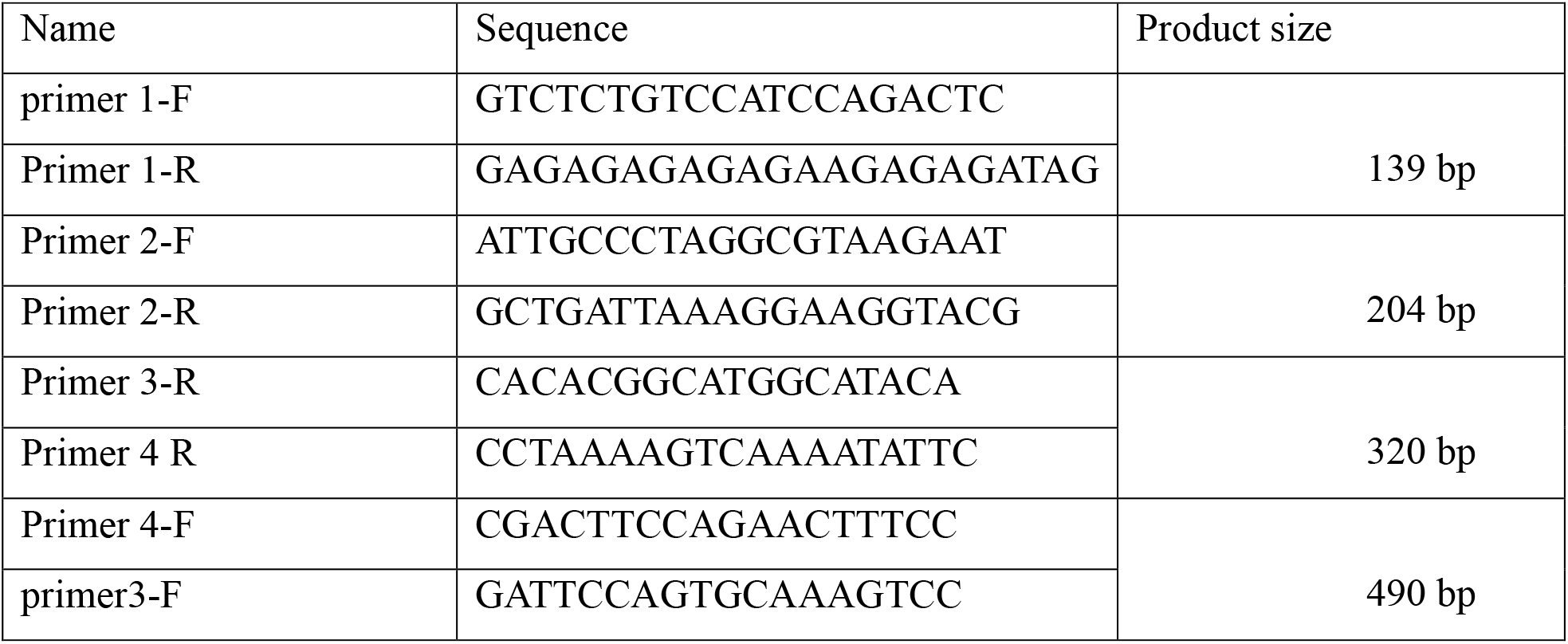
The Name, sequence and Product size of the Primer pairs used for the PCR purification.

For cloning applications, restriction digestion was performed on 400 ng of the pU6-pegRNA-GG-acceptor plasmid (Plasmid #132777 from Addgene)(Liu et al. 2022). The digestion reaction was carried out using the BsaI enzyme in a 10 µL reaction volume, incubated for three hours to ensure complete digestion. The digested plasmid was then purified using the same protocol described below for gel-extracted DNA, preparing it for ligation.

### DNA Purification from Agarose Gels and PCR Products

The protocol involves preparing a precipitation buffer, gel filtering tubes, and washing buffers. Specifically, the precipitation buffer was prepared by dissolving 15g of PEG (PEG6000-8000) and 8.77g of NaCl in approximately 30 mL of ultrapure water, adjusting the final volume to 50 mL. This viscous mixture required thorough mixing with a magnetic stir bar, with gentle heating if necessary to ensure complete dissolution. Optionally, filter sterilization was performed for specific applications **(Fig. 1)**.

**Figure 1:**
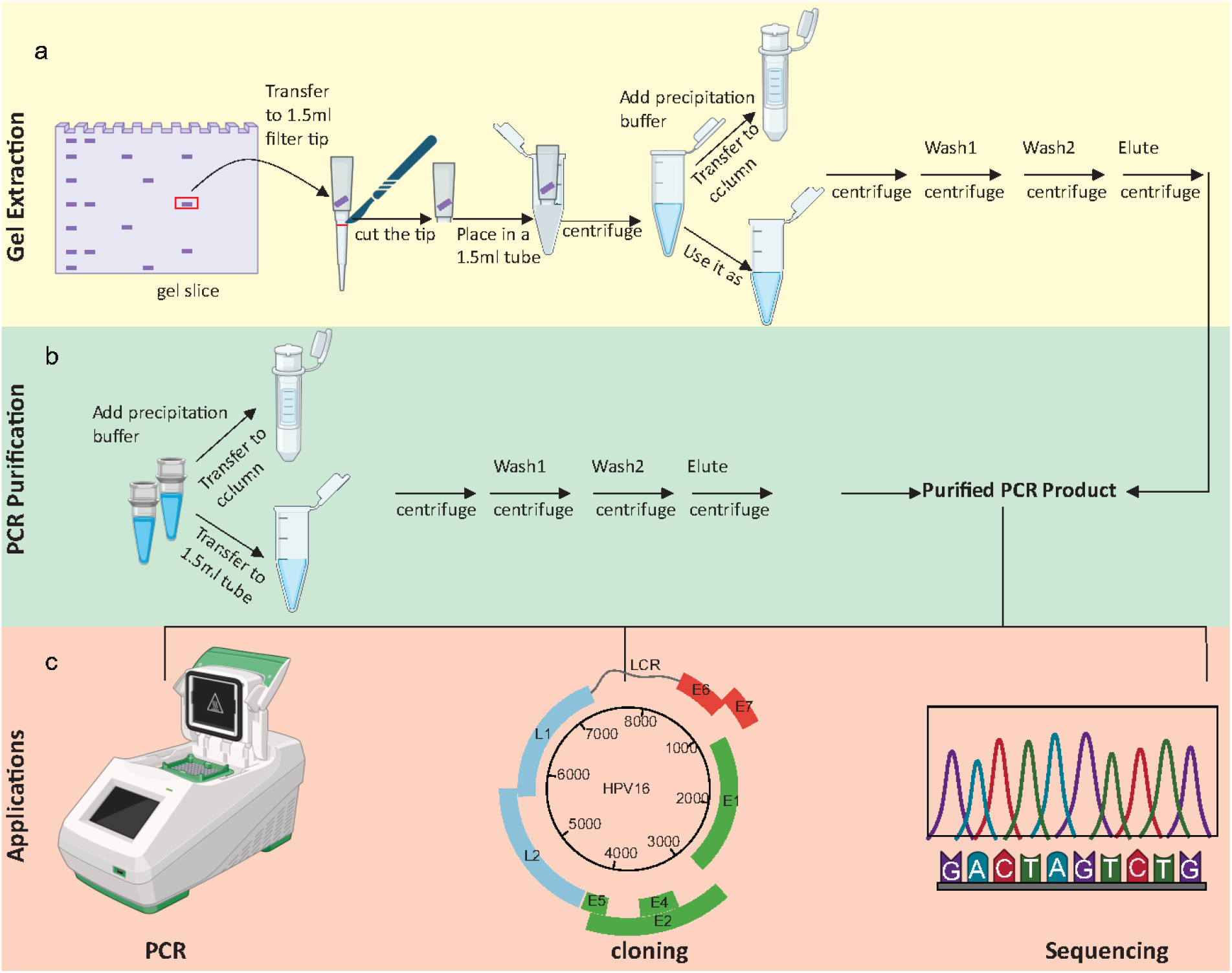
Workflow for purification. The process begins with gel extraction, where the desired gel slice is cut and transferred to a filter tube made as a result of cutting the blue filter (1.5ml) tip at the desired point. Following centrifugation, a precipitation buffer is added, and the mixture is transferred to a purification column. Subsequent washing steps (Wash 1 and Wash 2) are performed, and the final elution yields purified PCR products, ready for applications such as PCR cloning and sequencing.

The gel filtering tubes consisted of a 1 mL blue filter tip placed below the filter within a 1.5 mL microcentrifuge tube. A 50 µL PCR reaction setup was used in thin-walled 0.2 mL tubes for subsequent steps. For purification, 100% ethanol and 70% ethanol were prepared as washing buffers. After running the PCR product or restriction digestion on an agarose gel, the desired DNA band was excised and transferred into the filtering tube. Centrifugation at 10,000 rpm for 2 minutes allowed the DNA to elute, while the gel slice remained above the filter. Following elution, the precipitation buffer was added to the DNA sample, which was then centrifuged at 14,000 rpm for 5 minutes. Silica column purification was an alternative to direct centrifugation, though this option generally reduced DNA recovery.

The initial DNA pellet was washed with 100% ethanol, centrifuged again, and then subjected to a second wash with 70% ethanol. The purified DNA was resuspended in 20 µL of nuclease-free water or TE buffer, suitable for PCR amplification, molecular cloning, and sequencing. Quality and quantity were assessed using a spectrophotometer (Nanodrop). This same protocol was applied to PCR reactions, with DNA precipitation achieved by mixing the reaction with an equal volume of precipitation buffer and following the outlined centrifugation and elution steps.

### Statistical Analysis

Data from PCR amplification, DNA yield, and purity measurements were statistically analyzed to ensure reproducibility and assess the efficiency of the optimized protocol. Statistical analysis was performed using R, employing descriptive statistics (mean and standard deviation) to summarize DNA yield and purity.

## Results and Discussion

### 1. Comparison of Plasmid DNA and PCR Product Purification Methods

The comparison of different plasmid DNA and PCR purification methods, as illustrated in **(Fig. 2)**, evaluates the cost, time, concentration, and purity of DNA obtained using various approaches, including our optimized in-house protocol alongside Qiagen, Thermo Fisher, and WizBio commercial kits. This analysis provides insights into each method’s efficiency, feasibility, and potential applications in research settings. The left one is for the Plasmid and right side shows the PCR purification. In terms of cost, **Panel a** shows that the in-house method significantly reduces expenses compared to commercial kits, offering a notable financial advantage for laboratories with budget constraints. This cost-effectiveness could increase accessibility to high-quality DNA purification in resource-limited settings, promoting broader adoption of molecular techniques without the financial burden of proprietary reagents and equipment. **Panel b** highlights the time required for purification, with the in-house method demonstrating a marked reduction in processing time compared to the Qiagen and Thermo Fisher kits, which require longer durations. The efficiency of the in-house approach in saving time is particularly advantageous for laboratories aiming to increase throughput or streamline workflows, reducing delays between experiments and enabling more rapid sample processing(Lau et al. 2021). This is especially valuable in high-demand settings where rapid DNA purification is critical for timely project advancement(Tan and Yiap 2009). Furthermore, **Panel c** reveals that despite its lower cost, the in-house method achieves DNA concentrations competitive with those of the more expensive commercial kits. This finding supports the protocol’s suitability for applications where high DNA concentration is necessary, such as cloning or sequencing, and confirms its potential to meet the rigorous demands of molecular biology research without the added expense of commercial purification methods(Jun et al. 2012). Lastly, **Panel d** compares the purity of the DNA samples using the 260/280 absorbance ratio, a key indicator of DNA quality where values close to 1.8 reflect optimal purity with minimal protein contamination(Healey et al. 2014). The in-house method achieved a purity level comparable to the Qiagen, Thermo Fisher, and WizBio kits, demonstrating that it can reliably produce high-quality DNA suitable for downstream applications. This purity level is crucial for ensuring the consistency and reproducibility of results in sensitive molecular techniques, further validating the method’s utility in diverse research contexts(Mayjonade et al. 2016; Piepenburg et al. 2006). Overall, this analysis highlights the in-house DNA purification method as a cost-efficient, time-saving, and high-quality alternative to commercial kits. The benefits observed make it an attractive choice for laboratories with high-throughput needs, limited budgets, or resource constraints, offering a viable and accessible option to enhance molecular research capacity across various environments.

**Figure 2:**
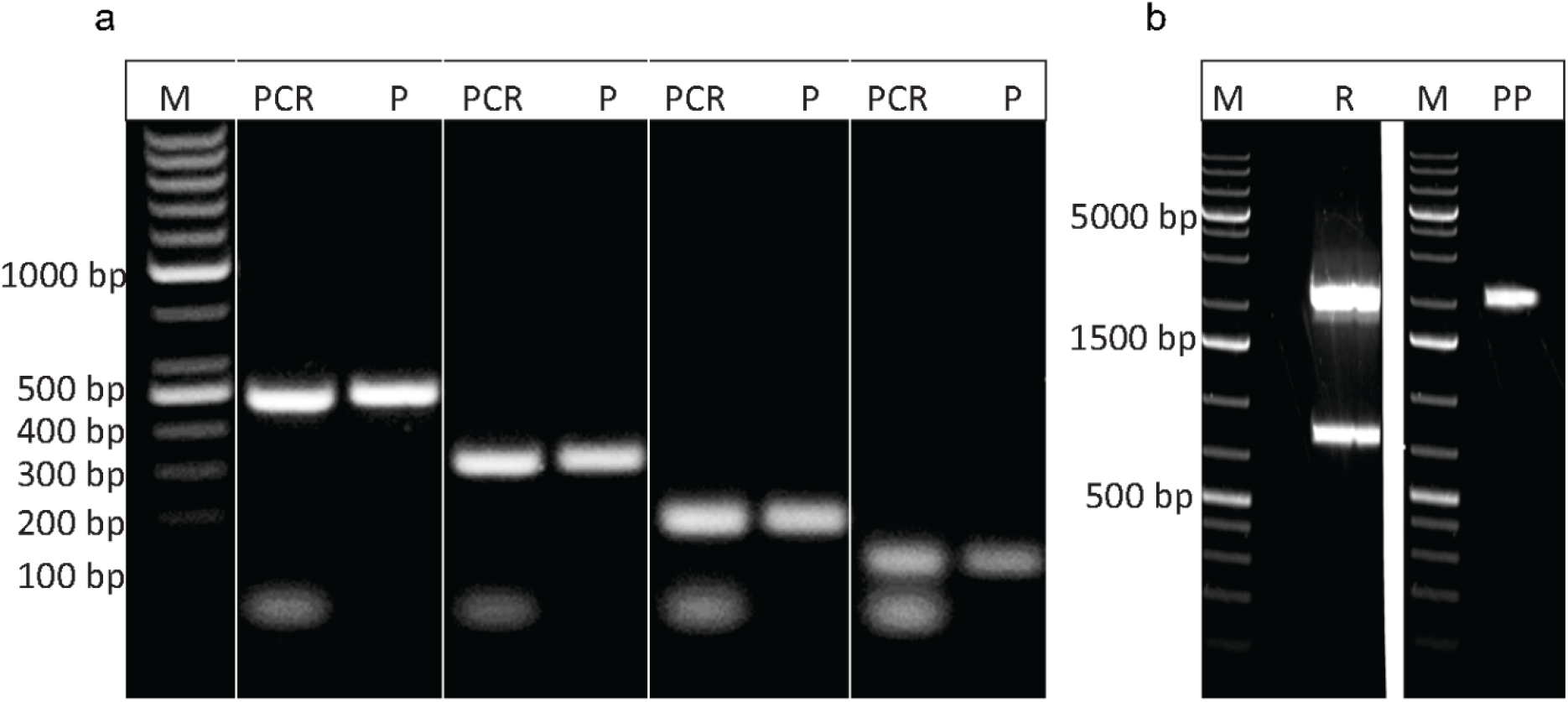
Comparison of plasmid (Right) and PCR (left) purification methods, assessing **(a)**cost (in USD), **(b)**time (in minutes), **(c)**DNA concentration (in ng/µL) with standard error, and **(d)**purity (260/280 ratio). Methods include in-house, Qiagen, Thermo Fisher, and WizBio, highlighting each approach’s relative efficiency, accessibility, and effectiveness in producing high-quality DNA suitable for downstream applications.

### 2. Restriction Digestion and Purification of Plasmid Samples

In our results, gel electrophoresis visually compares purified and unpurified PCR products, showcasing the efficacy of the in-house DNA purification protocol across a range of PCR amplicon sizes **(Fig. 3)**. The gel is structured into two panels, each emphasizing different aspects of the protocol’s effectiveness in enhancing DNA quality and clarity for various downstream applications. In **Panel a**, PCR products of specific sizes 139 bp, 204 bp, 320 bp and 490 bp are displayed, with each unpurified PCR product directly followed by its corresponding purified product (labeled “P”) in adjacent lanes. This arrangement of unpurified and purified products highlights the in-house protocol’s impact on band clarity and intensity, with purified samples showing significantly reduced background smearing and fewer non-specific bands than their unpurified counterparts. These improvements underscore the protocol’s effectiveness in producing clean PCR amplicons, which are critical for obtaining accurate and reliable results in molecular applications(Brennan et al. 2019). **Panel b** illustrates the restriction digestion of a plasmid sample, with the digested product labeled as “R.” The following lane, labeled “PP,” displays the purified version of this digested product. By comparing the restriction-digested plasmid before and after purification, **Panel b** further validates the protocol’s capacity to produce high-quality DNA that is suitable for sensitive downstream procedures, such as cloning, sequencing, or transformation(Whitlock et al. 2008). Additionally, a molecular marker lane (labeled “M”) is included in each panel to provide a reference for molecular weight, allowing for consistent size verification across purified and unpurified samples and thereby reinforcing the reliability of the purification results. Together, these findings presented in **Fig. 3** confirm that the in-house purification protocol yields DNA with reduced impurities and improved clarity, offering a viable and efficient alternative to commercial purification kits for laboratories requiring high-quality DNA for various applications.

**Figure 3:**
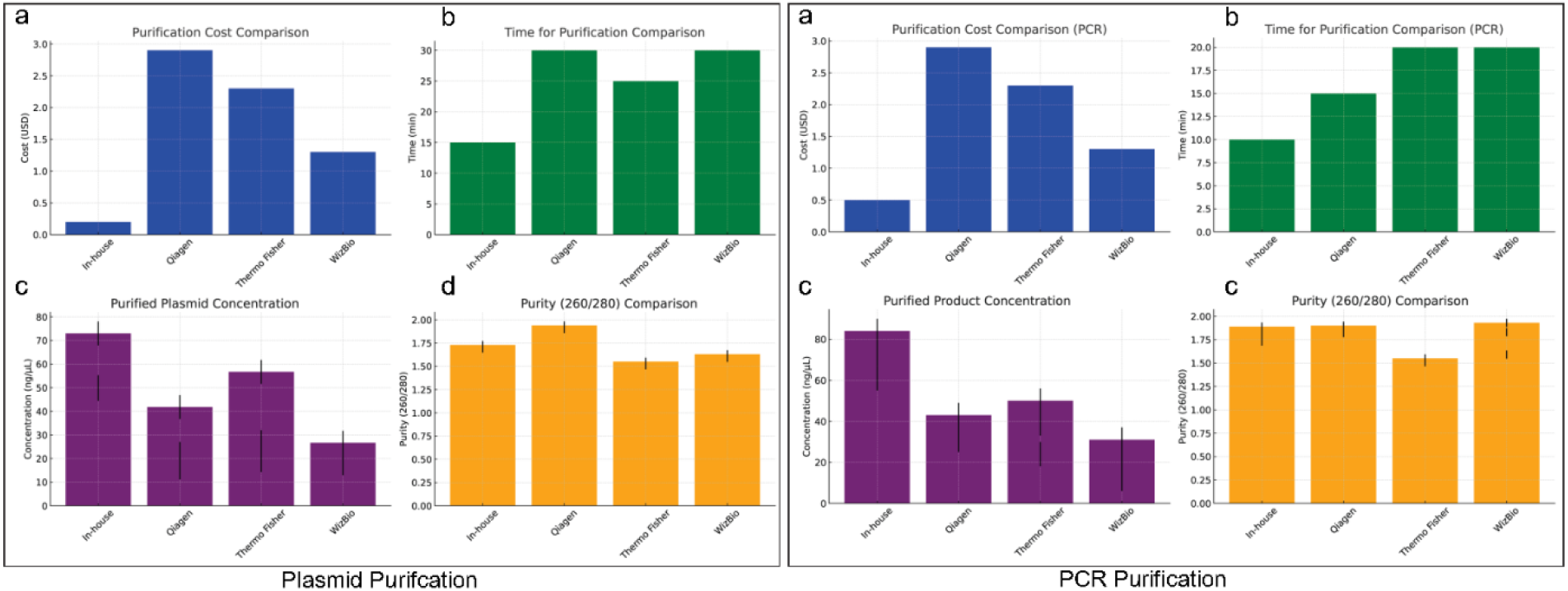
Gel electrophoresis comparison of purified and unpurified PCR products and plasmid samples. **(a)** Gel image of PCR amplicons of 139 bp, 204 bp, 320 bp and 490 bp, with each unpurified product followed by its purified form (labeled “P”). **(b)** Restriction-digested plasmid sample (labeled “R”) and its purified form (labeled “PP”), with the molecular marker 100 bp for PCR and 1kb for plasmid (labeled “M”) included for size reference across both panels.

### 3. Confirmation of Quality via Cloning and Sequencing

To assess the quality of the DNA purified from the gel using our in-house strategy, we performed pegRNA cloning through the Golden Gate assembly method, as outlined in **Fig. 4**. The pegRNA acceptor vector was linearized using the restriction enzyme BsaI, enabling quality selection through a color-based screening in E. coli. The disted plasmid was purified using inhouse methodology as previously described. Non-digested plasmids resulted in red colonies, while the addition of pegRNA protospacer, scaffold, and 3’ extension annealed oligonucleotides to the linearized vector, followed by T4 DNA ligase-mediated ligation, produced white colonies. This color distinction served as an initial screen for successful pegRNA integration, indicating high-quality, contamination-free DNA suitable for cloning. Subsequent confirmation of the pegRNA constructs was performed by PCR and Sanger sequencing, as shown in **Fig. 5**. These analyses verified the accuracy of the pegRNA insertion and the sequence integrity, affirming the effectiveness of the in-house purification method in yielding high-quality DNA appropriate for downstream applications.

**Figure 4:**
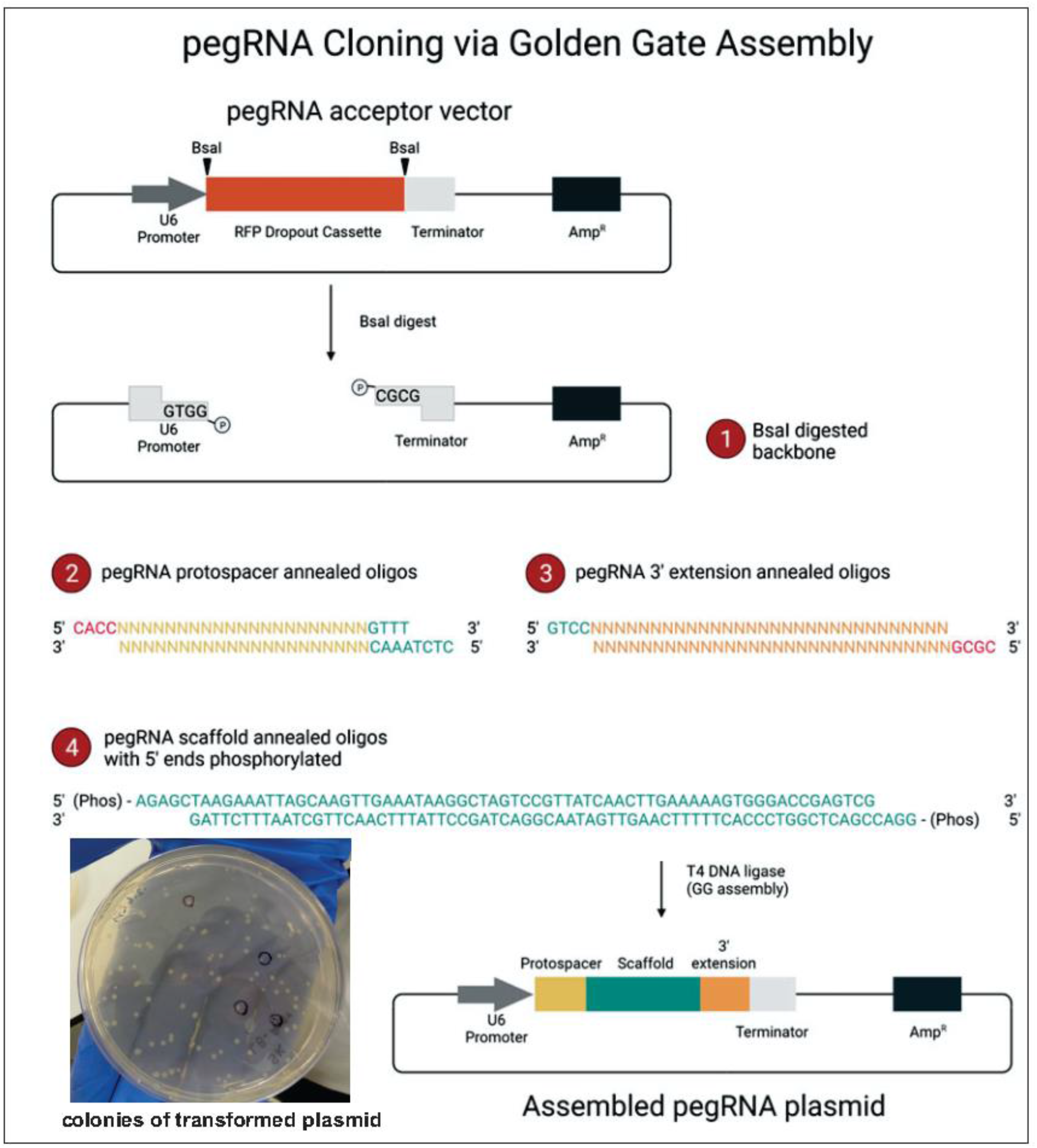
Schematic representation of the pegRNA cloning method using Golden Gate assembly. The pegRNA acceptor vector is first linearized by the restriction enzyme BsaI. The addition of the pegRNA protospacer, scaffold, and 3’ extension annealed oligonucleotides to the linearized vector, followed by ligation with T4 DNA ligase, results in successful assembly of the pegRNA construct. Transformed E. coli colonies on the plate display white color, indicating successful transformation and correct pegRNA assembly.

**Figure 5:**
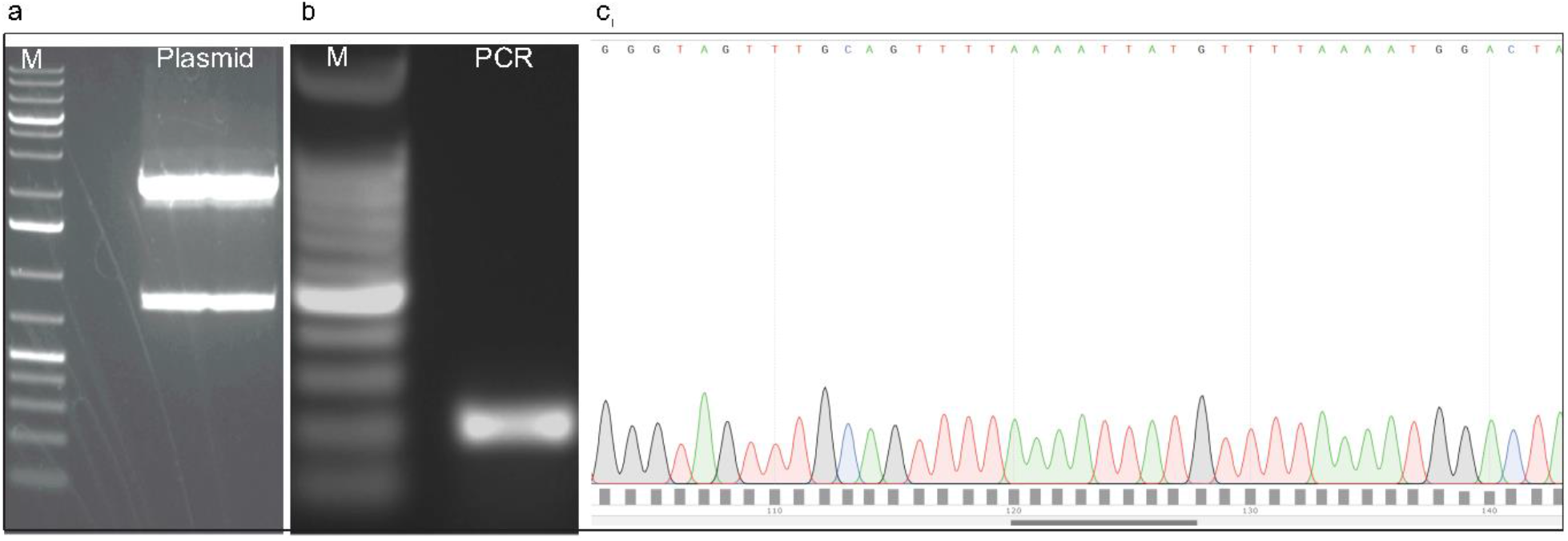
Confirmation of pegRNA constructs by PCR and Sanger sequencing. **(a)** Gel image showing the undigested plasmid following miniprep purification, with M representing the 1kb ladder (Thermo Scientific) and Plasmid indicating the miniprep-isolated plasmid. **(b)** Gel image showing successful PCR amplification from the miniprep plasmid, with M representing the 100bp ladder (Thermo Scientific) and PCR denoting the amplified product used for confirmation of transformation. (c) Sanger sequencing of the PCR product further validates the accuracy and integrity of the pegRNA construct, confirming its suitability for downstream applications. The combination of PCR amplification and Sanger sequencing demonstrates successful integration of the pegRNA sequence into the acceptor vector.

## Conclusion

This optimized, cost-effective DNA purification protocol offers a practical and efficient alternative to traditional commercial kits, particularly for laboratories operating under financial constraints. By utilizing a simple in-house method, it ensures high yields of sequencing-grade DNA from PCR products and agarose gels, suitable for various molecular biology applications. The substantial reduction in reagent costs, combined with faster processing times, makes this method ideal for high-throughput workflows. Its broad applicability and technical simplicity provide a valuable tool for researchers looking to maximize efficiency without compromising the quality of their results. This advancement addresses a critical need in molecular biology, particularly in resource-limited settings, and has the potential to significantly impact laboratory workflows globally.

## Acknowledgment

The authors would like to thank the staff and colleagues at the Centre for Regenerative Medicine and Stem Cells Research, The Aga Khan University, Karachi, Pakistan, for their invaluable support throughout the development and testing of this in-house DNA purification protocol.

## Author Contributions

SF designed and conducted the experiments, performed data analysis, and prepared the initial manuscript draft. MJ contributed to experimental design, data interpretation, and manuscript review. HZ assisted with experimental procedures, data collection, and processing. IH supervised the project, provided critical feedback on the methodology, and contributed to manuscript revision. AAM provided overall guidance, contributed to experimental design, and reviewed the final manuscript draft. All authors read and approved the final version of the manuscript.

## References

Ahmed N, Nawaz S, Iqbal A, Mubin M, Butt A, Lightfoot DA, Maekawa M (2013) Extraction of high-quality intact DNA from okra leaves despite their high content of mucilaginous acidic polysaccharides. Biosci Methods 4 (4):19–22

Aldrich J, Cullis CA (1993) RAPD analysis in flax: Optimization of yield and reproducibility using klen Taq 1 DNA polymerase, chelex 100, and gel purification of genomic DNA. Plant Molecular Biology Reporter 11 (2):128–141. doi:10.1007/BF02670471

Brennan S, Garcia-Castañeda M, Michelucci A, Sabha N, Malik S, Groom L, LaPierre LW, Dowling JJ, Dirksen RT (2019) Mouse model of severe recessive RYR1-related myopathy. Human Molecular Genetics 28 (18):3024–3036. doi:10.1093/hmg/ddz105

Carapito R, Bernardo SC, Pereira MM, Neves MC, Freire MG, Sousa F (2023) Multimodal ionic liquid-based chromatographic supports for an effective RNA purification. Separation and Purification Technology 315. doi:10.1016/j.seppur.2023.123676

Depristo MA, Banks E, Poplin R, Garimella KV, Maguire JR, Hartl C, Philippakis AA, Del Angel G, Rivas MA, Hanna M, McKenna A, Fennell TJ, Kernytsky AM, Sivachenko AY, Cibulskis K, Gabriel SB, Altshuler D, Daly MJ (2011) A framework for variation discovery and genotyping using next-generation DNA sequencing data. Nature Genetics 43 (5):491–501. doi:10.1038/ng.806

Diogo MM, Queiroz JA, Monteiro GA, Martins SAM, Ferreira GNM, Prazeres DMF (2000) Purification of a cystic fibrosis plasmid vector for gene therapy using hydrophobic interaction chromatography. Biotechnology and Bioengineering 68 (5):576–583. doi:10.1002/(SICI)1097-0290(20000605)68:5<576::AID-BIT13>3.0.CO;2-5

Ferreira P, Riscado M, Bernardo S, Freire MG, Faria JL, Tavares APM, Silva CG, Sousa F (2023) Pristine Multi-walled carbon nanotubes for a rapid and efficient plasmid DNA clarification. Separation and Purification Technology 320. doi:10.1016/j.seppur.2023.124224

Fu Y, Chen Q, Jia L (2021) RNase-free RNA removal and DNA purification by functionalized magnetic particles. Separation and Purification Technology 267. doi:10.1016/j.seppur.2021.118616

Healey A, Furtado A, Cooper T, Henry RJ (2014) Protocol: A simple method for extracting next-generation sequencing quality genomic DNA from recalcitrant plant species. Plant Methods 10 (1). doi:10.1186/1746-4811-10-21

Jun G, Flickinger M, Hetrick KN, Romm JM, Doheny KF, Abecasis GR, Boehnke M, Kang HM (2012) Detecting and estimating contamination of human DNA samples in sequencing and array-based genotype data. American Journal of Human Genetics 91 (5):839–848. doi:10.1016/j.ajhg.2012.09.004

Lau YL, Ismail IB, Mustapa NIB, Lai MY, Soh TST, Hassan AH, Peariasamy KM, Lee YL, Kahar MKBA, Chong J, Goh PP (2021) Development of a reverse transcription recombinase polymerase amplification assay for rapid and direct visual detection of Severe Acute Respiratory Syndrome Coronavirus 2 (SARS-CoV-2). PLoS ONE 16 (1 January). doi:10.1371/journal.pone.0245164

Liu G, Lin Q, Jin S, Gao C (2022) The CRISPR-Cas toolbox and gene editing technologies. Molecular Cell 82 (2):333–347. doi:10.1016/j.molcel.2021.12.002

Maurisse R, Fichou Y, De Semir D, Cheung J, Ferec C, Gruenert DC (2006) Gel purification of genomic DNA removes contaminating small DNA fragments interfering with polymerase chain reaction analysis of small fragment homologous replacement. Oligonucleotides 16 (4):375–386. doi:10.1089/oli.2006.16.375

Mayjonade B, Gouzy J, Donnadieu C, Pouilly N, Marande W, Callot C, Langlade N, Muños S (2016) Extraction of high-molecular-weight genomic DNA for long-read sequencing of single molecules. BioTechniques 61 (4):203–205. doi:10.2144/000114460

Piepenburg O, Williams CH, Stemple DL, Armes NA (2006) DNA detection using recombination proteins. PLoS Biology 4 (7):1115–1121. doi:10.1371/journal.pbio.0040204

Stadler J, Lemmens R, Nyhammar T (2004) Plasmid DNA purification. Journal of Gene Medicine 6 (SUPPL. 1):S54–S66. doi:10.1002/jgm.512

Tan SC, Yiap BC (2009) DNA, RNA, and protein extraction: The past and the present. Journal of Biomedicine and Biotechnology 2009. doi:10.1155/2009/574398

Vomelová I, Vaníčková Z, Šedo A (2009) Methods of RNA purification. All ways (should) lead to Rome. Folia Biologica 55 (6):243–251

Whitlock R, Hipperson H, Mannarelli M, Burke T (2008) A high-throughput protocol for extracting high-purity genomic DNA from plants and animals. Molecular Ecology Resources 8 (4):736–741. doi:10.1111/j.1755-0998.2007.02074.x

